# Structural heterogeneity of cellular K5/K14 filaments as revealed by cryo-electron microscopy

**DOI:** 10.1101/2021.05.12.442145

**Authors:** Miriam S. Weber, Matthias Eibauer, Suganya Sivagurunathan, Thomas M. Magin, Robert D. Goldman, Ohad Medalia

**Affiliations:** Department of Biochemistry, University of Zurich, Switzerland; Department of Cell and Developmental Biology, Northwestern University Feinberg School of Medicine, USA; Institute of Biology, University of Leipzig, Germany

**Keywords:** keratins, intermediate filaments, CRISPR knockout, cryo-electron microscopy, cryo-electron tomography

## Abstract

Keratin intermediate filaments are an essential and major component of the cytoskeleton in epithelial cells. They form a stable yet dynamic filamentous network extending from the nucleus to the cell periphery. Keratin filaments provide cellular resistance to mechanical stresses, ensure cell and tissue integrity in addition to regulatory functions. Mutations in keratin genes are related to a variety of epithelial tissue diseases that mostly affect skin and hair. Despite their importance, the molecular structure of keratin filaments remains largely unknown. In this study, we analyzed the structure of keratin 5/keratin 14 filaments within ghost keratinocytes by cryo-electron microscopy and cryo-electron tomography. By averaging a large number of keratin segments, we have gained insights into the helical architecture of the filaments. Interestingly, two-dimensional classification revealed profound variations in the diameter of keratin filaments and their subunit organization. Reconstitution of filaments of substantial length from keratin segments uncovered a high degree of internal heterogeneity along single filaments, which can contain regions of helical symmetry, regions with less symmetry and regions with significant diameter fluctuations. Cross section views of filaments revealed that keratins form hollow cylinders consisting of multiple protofilaments, with an electron dense core located in the center of the filament. These findings shed light on the complex architecture of keratin filaments, which demonstrate a remarkable degree of heterogeneity, suggesting that they are highly flexible, dynamic cytoskeletal structures.

## Introduction

Keratin Intermediate Filaments (KIFs) are an essential component of the cytoskeleton of epithelial cells. KIFs are classified as type I and type II Intermediate Filament (IF) proteins, according to their sequence (1-3). Keratins form a highly flexible and dynamic filamentous network in the cytoplasm (4-9). Their main known function is to protect the cell from external stresses by providing mechanical stability and ensuring the integrity of tissues through cell-cell and cell-matrix contacts. Point mutations in keratin genes are associated with cell and tissue instabilities and severe diseases, termed keratinopathies (3, 10). For example, the skin blistering disease Epidermolysis Bullosa Simplex (EBS) is caused by point mutations in the Keratin 5 (K5) and 14 (K14) genes (11-13). A key to understanding the function of KIFs in both normal and diseased cells is to unveil the structural organization of the filaments.

Keratin proteins are composed of three domains: a highly conserved α-helical central rod domain, known to facilitate filament assembly, and intrinsically disordered head and tail domains. The latter are highly post-translationally modified and are regarded as essential for regulatory functions and filament stability (14-16). Keratins are obligatory heterodimers, formed by the parallel and in-register assembly of an acidic type I and a basic type II keratin protein. Two dimers assemble into antiparallel tetramers, which can further align laterally and longitudinally to build mature keratin filaments (17-19). Longitudinally elongated tandem arrays of tetrameric subunits are termed protofilaments.

Although KIFs have been studied intensely for many years, details of their molecular architecture remain largely unknown (20). On the level of keratin monomers and dimers, crystallographic studies have provided high resolution insights into the organization of small regions of the central rod domain (18, 21-23) and molecular dynamics simulations have provided a 3D model of a complete K1/K10 dimer (24). Four different modes of tetramer assembly have been identified by cross-linking studies, describing how higher order oligomers form during filament assembly (17). It is expected that filament assembly starts by the formation of A_11_ tetramers, where the 1B domains of the rod of two adjacent dimers interact in a half-staggered, antiparallel arrangement (18, 19). Tetramers can elongate longitudinally by formation of A_22_ interactions, where the 2B domains of adjacent rods overlap (18, 19). Laterally, neighboring tetramers interact via A_12_ bindings to form 10 nm wide filaments (18, 19). However, little is known about the 3D high-resolution structure of mature keratin filaments. It is generally accepted that keratin filaments are helical assemblies consisting of multiple protofilaments (25-27), which form a cylindrical tube (15). However, the exact number of protofilaments per filament and therefore the number of keratin monomers per cross section is still debated and may vary with respect to the specific type I/type II keratin pairs expressed. Mass-Per-unit-Length (MPL) measurements of recombinant *in vitro* assembled K8/K18 filaments have suggested that they are built from 16 – 25 monomeric protein chains in cross section, depending on the ionic strength of the buffer and the assembly time (28). Interestingly, MPL measurements of KIFs assembled *in vitro* from keratins extracted from human epidermis have revealed that the majority of them contain 13 – 16 polypeptides in cross section, with fewer numbers of KIF comprised of either 20 – 26 or 28 – 35 polypeptides (29). These variations have been attributed to structural polymorphism of KIFs and apparently occur by varying the number of protofilaments along and among keratin filaments (29). In addition, there are conflicting views as to whether KIFs are hollow or filled tubes and whether they contain an internal structure (15, 30-34).

Here, we studied the structure of native cellular K5/K14 filaments by Cryo-Electron Microscopy (cryo-EM) and Cryo-Electron Tomography (cryo-ET) (35). Since the expression of keratin isoforms is variable and complex in cultured cells, we prepared a cell line expressing filaments composed of K5/K14 only and studied their architecture within cells that were grown and lysed on EM grids, i.e. ghost cells. This process avoids potential structural artifacts due to *in vitro* assembly of KIFs. Our cryo-EM analysis revealed the remarkably heterogenic nature of keratin filaments and uncovered changes in the diameter and the helical pattern propagating along the filament. Cryo-ET and analysis of filament cross sections revealed that the K5/K14 filaments are composed of a hollow cylinder with an internal electron dense core. The wall of the cylinder is constructed of a ring of six protofilaments. Our results quantify the flexibility of keratin filaments and uncover the immense structural heterogeneity of individual K5/K14 filaments.

## Results

### Generation of mouse keratinocytes expressing only K5/K14 filaments

Heterogeneity is a challenging problem that hampers structural determination (36). Thus, the occurrence of multiple keratin pairs in most epithelial cells hinders the structural analysis of keratin filaments in their native environment, due to intrinsic structural heterogeneity (37). In this study, we set to gain insights into the architecture of cellular K5/K14 filaments. Therefore, we utilized the murine keratinocyte cell line KtyI^-/-^ K14, which expresses the K5, K6 and K14 proteins as their only keratin isoforms (38). To reduce the keratin expression to K5 and K14 only, CRISPR/Cas9 was used to knock out the K6a and the K6b gene, which share 92.6 % sequence identity (Material and Methods section). Although a small amount of wild-type K6b DNA was retained (Figure 1–figure supplement 1A, B), immunostaining revealed that no filaments containing K6 assemble in the resulting K5/14_1 cell line, and therefore do not affect the structural analysis carried out in our study (Figure 1–figure supplement 1C). We therefore conclude that the keratin filaments in this cell line consist only of K5/K14 protein pairs. A careful analysis of the KIF network after the CRISPR/Cas9 knockout procedure by confocal fluorescence microscopy indicated no obvious impact on the K5/K14 filament network (Figure 1A, Figure 1–figure supplement 1C).

**Figure 1.**
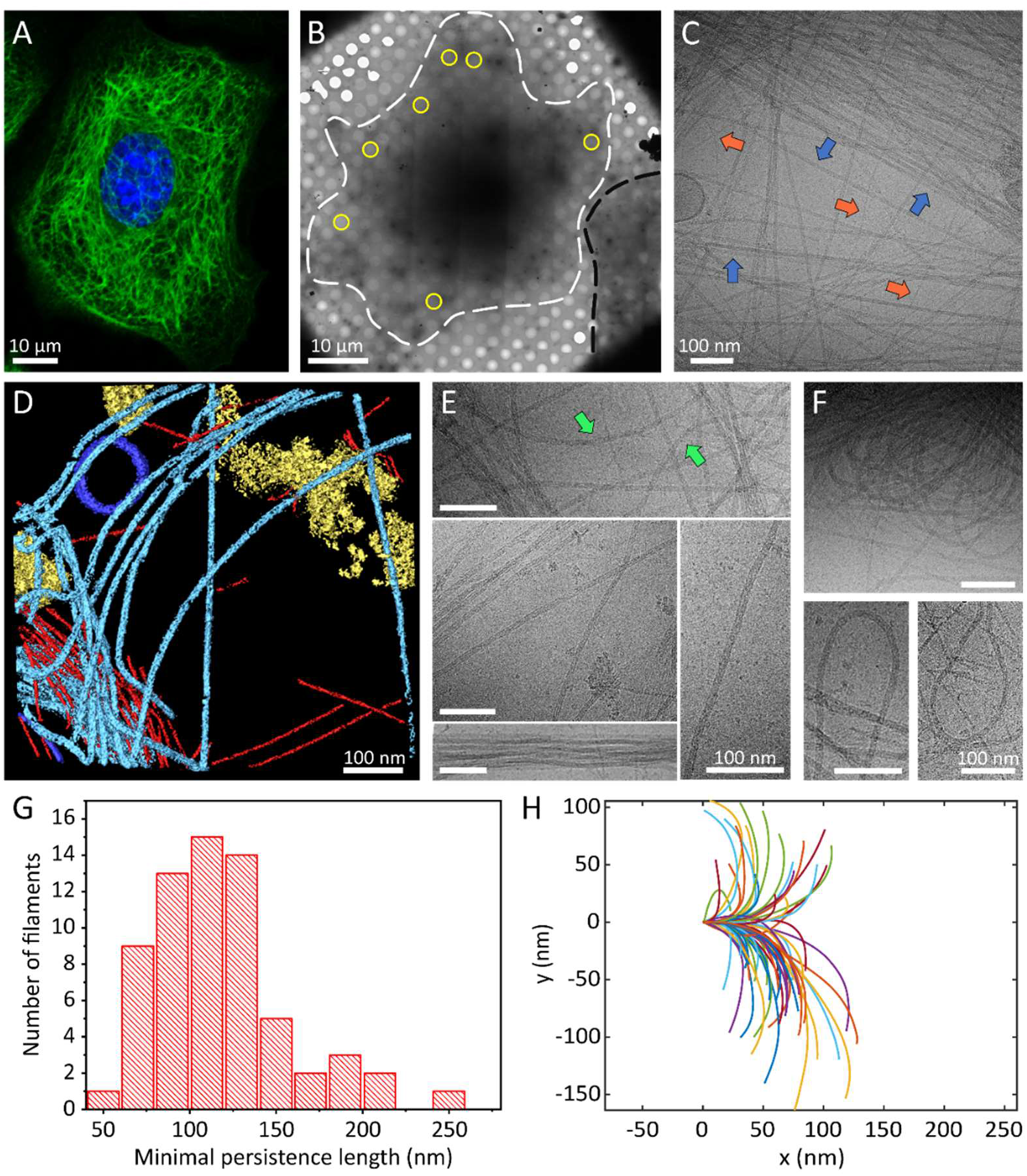
Cellular K5/K14 filaments as revealed by light and cryo-electron microscopy. (A) The murine keratinocyte cell line K5/K14_1 expressing only K5 and K14 filaments forms a complex KIF meshwork, as revealed by confocal immunofluorescence. Cells were stained for K14 (green) and chromatin (blue). (B) Ghost cells were analyzed by cryo-EM and cryo-ET. Low magnification image of a cell grown on an EM-grid and treated with detergent prior to vitrification. Cell boundaries (dashed white line) are detected as well as a neighboring cell (dashed black line). Typical regions that were analyzed by cryo-EM are marked (yellow circles). (C) A typical cryo-EM micrograph of a ghost cell imaged at a higher magnification allows the detection of keratin filaments and other cytoskeletal elements. Keratin filaments (blue arrows) and actin filaments (orange arrows) are distinguished by their characteristic diameter. A large keratin bundle is visible in the top right corner. (D) Surface rendering view of a cryo-tomogram of a ghost cell. Keratin filaments (light blue), actin filaments (red), vesicles (dark blue) and cellular debris (yellow) were manually segmented. (E) Different organizations of keratin filaments observed in the electron micrographs, including straight filaments (middle), curved (top, green arrows) and bundled filaments (bottom left). Scale bars: 100 nm. (F) Highly bent keratin filaments are found within ghost cells. Scale bars: 100 nm. (G) Quantification of the minimal apparent persistence length measurements performed on 65 highly bent keratin filaments. (H) A plot combining 65 contours of filaments that were used for the minimal apparent persistence length measurements in (G). Individual filaments, shown in different colors, are aligned at their origins for visualization purposes.

### Cellular keratin filaments revealed by cryo-electron microscopy

K5/14_1 cells were cultured on cryo-EM grids and subjected to cytoskeleton extraction buffer that permeabilizes the cells and removes soluble cytoplasmic components and nuclear structures, producing IF enriched ghost cells (Material and Methods section) (39-44). The ghost cells were instantly plunge frozen and imaged by cryo-EM and cryo-ET (Figure 1B-D). Keratin filaments could be easily identified in cryo-EM micrographs (Figure 1C, blue arrows, Figure 1–figure supplement 1D), while the 3D organization of the keratins within the ghost cells was revealed by cryo-ET (Figure 1D light blue). Actin filaments were detected in the sample as well (Figure 1C, orange arrows and D, red, Figure 1– figure supplement 1D) and used as an internal quality control for structural preservation by the extraction protocol and for cryo-EM image quality. Under these conditions, the structure of cellular F-actin was resolved to 6.1 Å (Figure 1–figure supplement 2).

Keratins form a complex filamentous network, including thick bundles containing numerous filaments, meshworks, in which filaments are often crossing and interacting with each other, as well as long stretches of individual filaments (Figure 1C, E, Figure 1–figure supplement 1D). Keratin filaments exhibit a wide range of shapes suggesting a high degree of flexibility. While some filaments are straight over long distances (several hundreds of nm), others exhibit a wavy appearance (Figure 1E, arrow). Additionally, highly bent keratin filaments are frequently detected (Figure 1F, Figure 1–figure supplement 1D). We traced 65 of these highly bent filaments and determined their minimal apparent persistence length (contour length upon 90° turn, Figure 1H). Keratin 5/14 filaments are able to undergo a 90° turn within 118.4 ± 39.2 nm (Figure 1G), similar to distances observed for nuclear lamins (40, 45). These measurements were conducted only on the sub-population of highly bent filaments and not on the full range of shapes detected (e.g. straight filaments), as their varying behavior prohibited us to describe all of them with a single measure (46). Interestingly, some filaments undertake even 180° turns within a few hundred nanometers without breaking and change directions multiple times within the inspected field of view (Figure 1F). In agreement with previous *in vitro* assembled keratin analyses, the remarkable flexibility of keratin filaments supports their ability to maintain filament integrity even under extreme conditions (47-49).

### Heterogeneity in filament diameter and helical pattern

To obtain deeper insights into the architecture of keratin filaments, we extracted and structurally averaged 55 nm long straight keratin segments picked along filaments in ∼1700 cryo-EM micrographs (50, 51). A helical pattern that spirals along the long filament axis can be detected in several class averaged structures (Figure 2A, Figure 2–figure supplement 1A). While several structural classes show a clear helical pattern, others reveal elongated, rather straight sub-structures without an apparent helical symmetry (Figure 2A, bottom, Figure 2–figure supplement 1A). Transition regions between the two distinguished patterns can also be detected (Figure 2A, arrows). The mean diameter of keratin filaments, as determined by direct measurement of intensity line-profiles through the class averages, is 10.1 ± 0.5 nm (Figure 2B, C), in agreement with previous observations (19). A mean intensity line-profile through a lateral average of the most populated classes defined the edges of the filaments as well as a central density peak (Figure 2C). The outer boundaries of the filaments show the highest electron density values and therefore are their most pronounced structural features (i.e., the filament diameter), while a central density peak with slightly lower intensity is also apparent. This analysis further revealed a subset of structural classes with a much larger diameter than the majority of filaments (Figure 2D). Intensity line profiles of a thicker class (black asterisk) and a more frequently detected class (blue asterisk) indicate a 30% difference in filament diameter, 13.2 nm vs 10.1 nm, respectively (Figure 2E). Moreover, the internal structure of the thicker classes diverges from the classes shown in Figure 2A. Specifically, some classes reveal two distinct linear electron densities within the filament (Figure 2D, arrowheads), indicating a less dense packing of the individual protofilaments as compared to the compact classes. Others capture transitions between a thinner and a thicker region along an individual filament (Figure 2D, arrows). These findings indicate deviations in the organization of protofilaments, reflecting structural heterogeneity along individual filaments.

**Figure 2.**
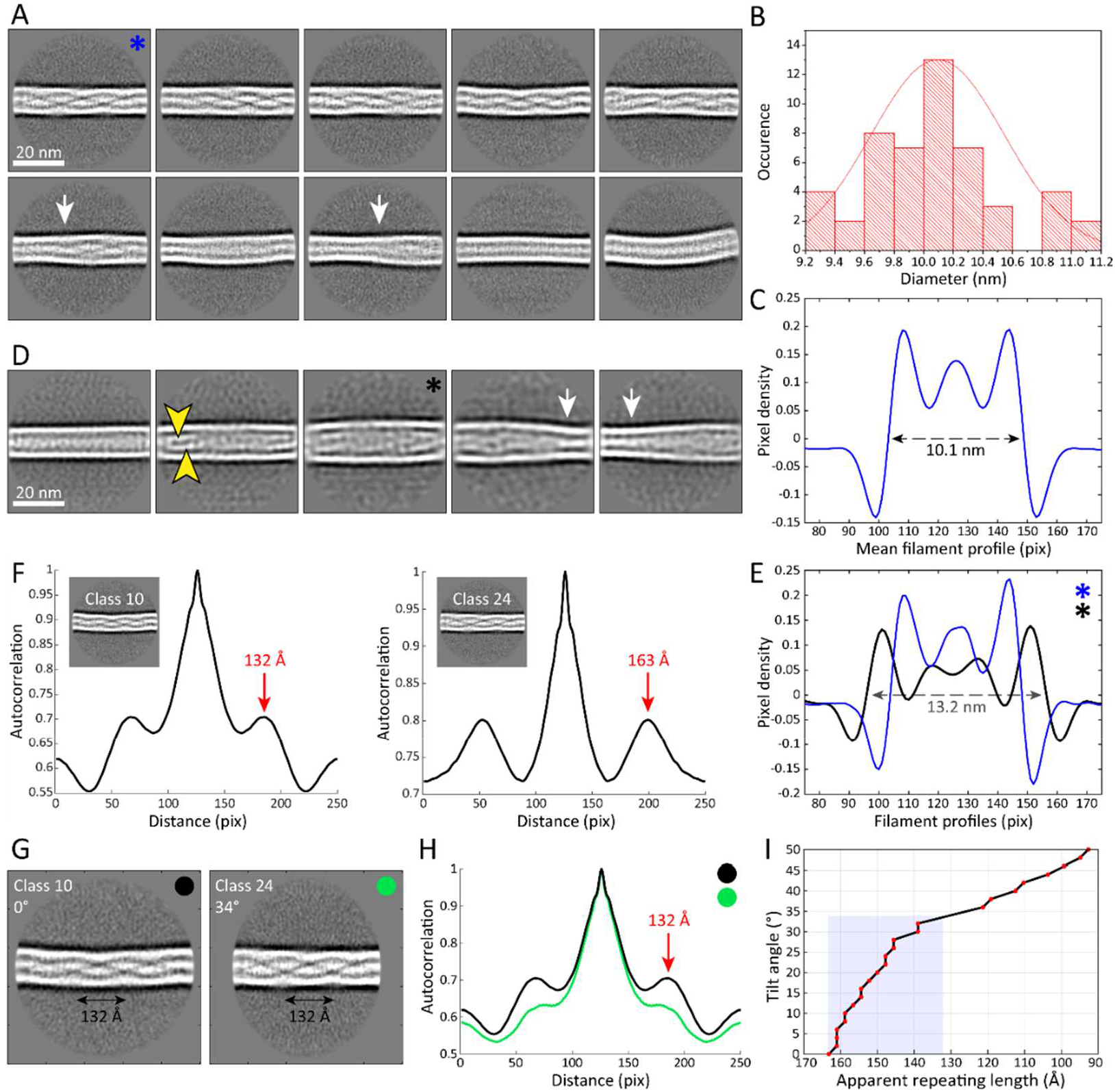
The architecture and heterogeneity of keratin filaments. (A) Ten of the most populated 2D class averages of keratin segments. High electron density is shown in white. Arrows indicate transition regions between helical and straight-line patterns. (B) Distribution of filament diameters as measured in 50 2D class averages (Figure 2–figure supplement 1A). (C) Mean intensity line-profile through all classes used in (B). The mean filament diameter (10.1 nm) is indicated. (D) Subset of keratin class averages showing larger filament diameters. Two individual filamentous densities are often detected within a filament (yellow arrowheads). Additionally, transition regions between thinner and thicker filament regions are detected (white arrows). (E) Intensity line-profiles through a narrow and a wide class indicated by blue and black asterisk (in (A) and (D)), respectively. Diameters of 10.1 nm and 13.2 nm (arrow) were detected. (F) Autocorrelation spectra of the displayed keratin classes (insets). Peaks of the autocorrelation function corresponding to the distance between repetitive elements along the filament are indicated (arrows). (G) To show that out-of-plane tilting of KIFs can shift the autocorrelation peaks, Class 24 was tilted *in silico* by 34°, while Class 10 is untilted. The apparent repeating distance of both classes is indicated. (H) Autocorrelation spectra of the classes shown in (G). After tilting of Class 24 by 34°, both classes show an autocorrelation peak at the same marked position, an indicator that filament tilting might be the reason for the different repeat distances observed in the 2D classes. Green and black dots indicate which curve belongs to which class in (G). (I) Dependence of the apparent repeating length (autocorrelation peaks) on the filament tilt angle, measured by tilting Class 24 from 0° to 50° and calculating corresponding autocorrelation spectra. The gray area indicates the range of repeat distances found in keratin 2D classes.

In order to determine the repeat distances of the helical patterns observed in the class averages we calculated autocorrelation spectra for each helical class (Figure 2F, Figure 2–figure supplement 1D). Using this approach, the repeating units along the length of the filament can be determined (52). We found that the repeat distance of the helical pattern varies dramatically among different classes, ranging from ∼132 Å to ∼163 Å (Figure 2F). Between these two extremes numerous distinct values for the repeat distance of the helical pattern can also be identified (Figure 2–figure supplement 1D). Since keratin filaments are very flexible and they form a 3D network in cells, we exploited the possibility that the different repeat distances reflect filaments that are oriented out of plane. The projection of a tilted filament in our cryo-EM micrographs would therefore yield classes with an apparent shorter repeating pattern. In this case, the class with the longest repeating distance would reflect the untilted filament, while all other repeating patterns would originate from different degrees of tilting. Based upon this reasoning, we tilted and projected the class with a repeat distance of 163 Å *in silico* and retrieved similar repeat distances as seen in the real classes (Figure 2 I). We showed that a tilt of up to ∼34° can induce shortening of the repeating pattern from 163 Å to 132 Å (Figure 2 G, H). Therefore, tilting between 0° - 34° would explain the variations that were detected in the repeating pattern of the keratin classes (Figure 2 I). With ice thicknesses of up to ∼300 nm and a minimal apparent persistence length of ∼118 nm, this amount of tilting can be expected and further analysis by cryo-ET revealed that even higher degrees of tilting are possible (see below).

### Reconstituting keratin filaments from the class averages

The 2D class averages allowed us to identify structural differences in 55 nm long keratin segments. To understand how these structural features are organized at the level of long keratin filaments, it was important to determine how the class averages are arranged along keratin filaments which are up to several hundreds of nanometers in length. For this purpose, we utilized a back-mapping strategy that permits the reconstitution of the original filament out of 2D class averages (Figure 2–figure supplement 1A, B) (41, 53). Therefore, every segment was represented by its corresponding 2D class image, which was inversely transformed, so that it matched the original orientation of the raw segment. Then it was plotted at the original coordinate position, where the raw segment was selected from the electron micrographs. In this fashion, we assembled the original keratin filaments made out of the respective 2D class averages (Figure 3–figure supplement 1A), which were subsequently extracted and straightened. Since these reconstituted filaments are assembled from class averages, their signal-to-noise ratio is drastically improved compared to the raw filaments. This approach allowed us to study long stretches of keratin filaments with improved resolution up to ∼12 Å (Figure 3, Figure 2–figure supplement 1C).

**Figure 3.**
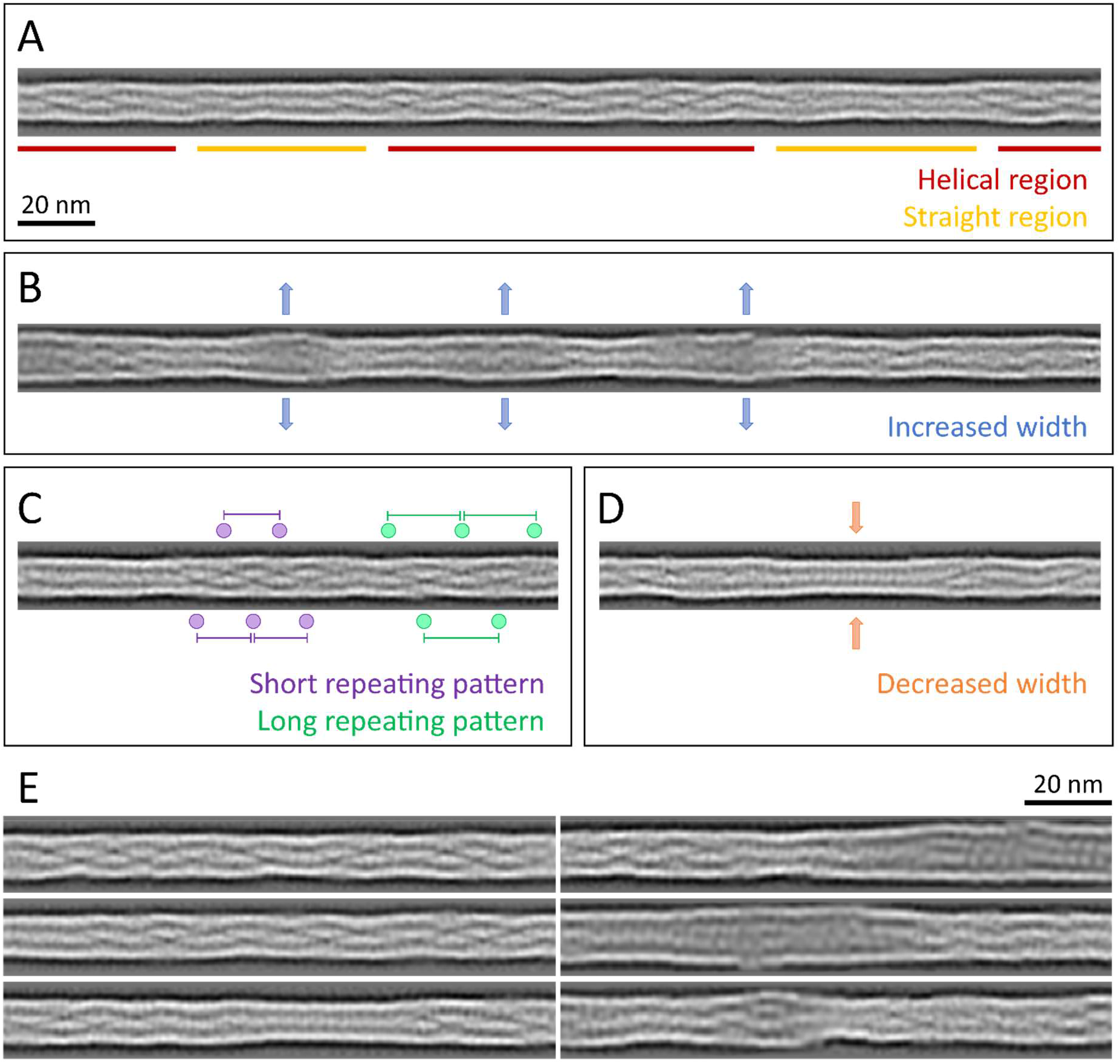
Structural polymorphism along keratin filaments. Reconstituted filaments provide a realistic view of the KIFs at higher resolution (see Materials and Methods). (A) – (D) Scale bar: 20 nm. (A) A typical keratin filament consisting of various regions with helical and straight-line patterns, indicated by red and yellow lines, respectively. (B) Keratin filament displaying diameter fluctuations. Areas of increased diameter are indicated (blue arrows). (C) Keratin filament showing helical patterns exhibiting different repeat distances, indicating modulations within the ice layer. Short repeating patterns (purple) indicate higher tilt angles in comparison to longer repeating patterns of in-plane filament stretches (green). (D) Keratin filament displaying diameter fluctuations. Areas of decreased diameter are indicated (orange arrows). (E) A collage of six reconstituted keratin filaments showing structural diversity.

The appearance of these reconstituted keratin filaments is very heterogenous (Figure 3, Figure 3– figure supplement 1). Overall, they consist of patches of helical regions with clear repetitive patterns (Figure 3A, red, E), which are frequently interrupted by straight patterned stretches with less defined features (Figure 3A, yellow, E). The helical as well as the straight-line stretches are variable in length and frequency. While some filaments consist of mostly helical stretches, others are mixed or exhibit a mostly straight-line appearance (Figure 3E, Figure 3–figure supplement 1B). Additionally, the diameter

fluctuates along a single filament (Figure 3B, D). For example, KIFs have regions of increased width up to 13.2 nm that often allow the identification of individual sub-chains (Figure 3B, E), as well as thinner regions with widths down to 9.2 nm (Figure 3D, E). The thinner regions usually display a straight pattern, whereas not all straight regions show a decrease in diameter. Interestingly, the reconstituted filaments revealed that helical regions with different repeat distances, identified in the 2D class averages (Figure 2), can co-exist along a single filament (Figure 3C). This indicates that individual filaments changed their tilt angle along the course of the filament and ran through different z-heights of the ghost cell volume. Helical patterns with different repeat distances, indicating different tilt angles, lie in close proximity along KIFs, where they appear to transition smoothly into each other. These structural transitions reveal that keratin filaments constantly fluctuate in the z-direction and thus appear to be as flexible in the z-direction as they are in the *xy* plane.

Overall, reconstituted keratin filaments reveal an enormous amount of structural heterogeneity (Figure 3E, Figure 3–figure supplement 1C). Every filament examined displays a unique phenotype, which demonstrates that keratin filaments are as versatile as the challenges they encounter in a living cell.

### Keratin filaments are hollow cylinders with an internal electron dense core

A careful analysis of our dataset revealed several cryo-EM micrographs and multiple cryo-tomograms (21 out of 44) which contain KIFs that undergo a 90° turn along the thickness of the sample and therefore allow the observation of direct cross sections of the keratin filaments. This behavior is quite remarkable, as it was never seen in tomograms (n = 225) of cellular vimentin intermediate filaments, imaged within detergent extracted mouse embryonic fibroblasts (MEF) (unpublished, Figure 4–figure supplement 1A, B). Analysis of cross sections revealed that keratin filaments are hollow cylinders, in which an internal electron dense core is found (Figure 4A-E). This finding agrees with previous studies, which predicted that keratins contain internal mass, but less than anticipated for a completely filled filament (15, 29-34). Moreover, we could identify individual sub-filaments, which form a hexameric ring structure in cross section (Figure 4B). Based on this geometry and previous literature, it is likely that the sub-structures represent tetrameric protofilaments and therefore the mature filament would be composed of ∼6 protofilaments to yield ∼24 polypeptides in cross section (15). This agrees with previous mass-per-unit-length analyses of epidermal keratins and keratins from simple epithelia (28, 29). Depending on the individual filament and tomographic slice, the number of visible protofilaments varies, which might indicate polymorphism, i.e. a variable number of protofilaments building the KIF. However, it might also be an imaging artifact. When following a filament in cross section through the tomographic volume, in certain slices densities of neighboring protofilaments seem to merge into one continuous structure, indicating that there are positions along the filament where the protofilaments are interacting so tightly that they could not be resolved individually (Figure 4E, Figure 4–figure supplement 1C). Other positions reveal more than six protofilaments, which may reflect overlap regions between tetramers along the keratin filaments.

**Figure 4.**
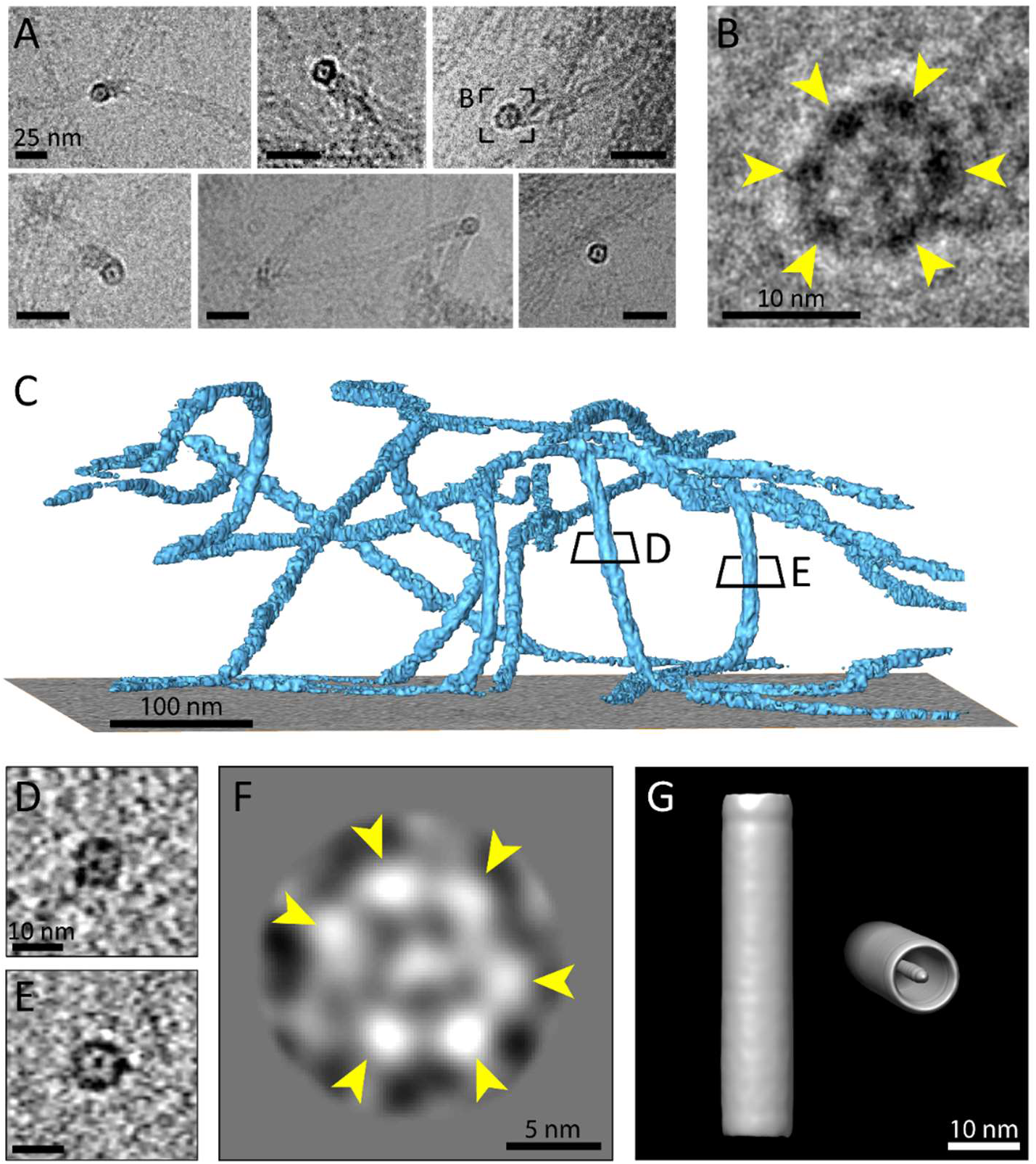
Multiple protofilaments and an internal electron dense core are canonical components of keratin filaments. (A) Cross section views of keratin filaments detected within the cryo-EM micrographs of ghost cells. An electron dense core is visible in the center of the keratin tube. Scale bars: 25 nm. (B) Zoomed-in view of the area boxed in (A). The cross section view reveals an internal core surrounded by six protofilaments as constituents of the tube (yellow arrowheads). (C) A surface rendered tomogram of a ghost cell was rotated in order to show the modulation of the keratin filaments within the ice layer. The three-dimensional keratin network is visualized (light blue). The level of the support is shown as a gray colored slice. Tomographic slices through vertically oriented filaments showing cross section views are indicated by boxes. (D) - (E) 7 nm thick xy-slices of the areas indicated in (C), showing KIFs as tube-like structures with a central density. Individual protofilaments can be identified. Scale bars: 10 nm. (F) A 2D class averaged structure of cross section views extracted from 19 individual regions of vertically oriented filaments, revealing the six individual protofilaments constituting the keratin filament tube (yellow arrowheads). (G) Low resolution 3D model indicating the overall dimensions of a keratin filament and the presence of the central density. The structure was calculated template-free by randomizing the rotation angle of extracted keratin segments. Left: Side view. Right: Tilted cross section view revealing internal electron dense core.

To study the number of protofilaments in more detail, we selected 710 cross-sectional views of filaments found within the cryo-tomograms and subjected them to 2D classification, revealing a symmetric hexameric class (Figure 4F). Six individual protofilaments can be clearly distinguished in the ring (arrowheads), which agrees with our studies of the raw data. As expected, additional classes were found showing deviations from this hexameric structure (Figure 4–figure supplement 1D). These structural differences may represent actual changes in symmetry or deviations from a perfect perpendicular cross section.

Finally, to get an impression of a mature keratin filament in three dimensions, we generated a low-resolution 3D model of a keratin filament from our dataset of 55 nm long segments using the Relion software package (50, 51). By randomizing the rotational angle along the filament axis, a template-free unbiased model was generated (Figure 4G). The 3D model strengthens our findings that keratin filaments are formed as hollow cylinders with a central electron density. Due to randomization of the rotation angle, the individual protofilaments are not resolved in this structure, however, it provides a view of the central density and the overall dimensions of the K5/14 intermediate filaments.

## Discussion

Keratin intermediate filaments are major components of the cytoskeleton that are involved in many cellular processes (54-56). However, due to their flexibility, heterogeneity and yet to be resolved symmetry, a high-resolution structure of keratin filaments in their native state has not been obtained to date. In this study, we describe novel insights into the architecture of *in vivo* assembled K5/K14 filaments by imaging them directly within ghost cells. This approach has enabled us to study native KIFs containing all their post-translational modifications, which are known to play an important role in their assembly and function (57). Moreover, this approach circumvents the need of denaturing and renaturing the proteins prior to *in vitro* assembly, a step which likely increases structural polymorphism (15).

The K5/K14 filaments were detected in ghost cells as individual, separated filaments or in bundles. In this study we have focused on individual filaments for technical reasons, as KIF bundles are dense, highly complex structures and would be unsuited for our averaging procedures (58). Individual filaments were found to be very flexible and showed a high degree of bending within a few hundred nanometers. We determined the minimal apparent persistence length of highly bent KIFs to be 118.4 ± 39.2 nm. Thus, intact keratin filaments can undergo a 90° turn within a distance of 2 – 3 dimer lengths (dimer length ∼44 nm (24, 59)). When compared to microtubules or actin filaments, which have persistence lengths of ∼7 – 22 µm and several mm, respectively, keratin filaments are much more flexible (60). Their long α-helical building blocks allow them to accommodate high bending, similar to nuclear lamins (40, 61). Previous studies showed that the persistence length of K8/K18 filaments can vary significantly between 300 – 650 nm depending on the method used (46, 62, 63). However, within the cell, individual filaments are incorporated into a network and therefore are not in a relaxed equilibrium state as expected *in vitro*. Thus, we suggest ∼118 nm as the minimal apparent persistence length of the K5/K14 sub-population of highly bent filaments.

The high flexibility and modulation of keratin filament orientation is also apparent as filaments can span through the entire thickness of the ice on a cryo-EM grid. The filaments are often tilted out of the *xy* plane and form a wavy network in all three dimensions. Cryo-tomograms of ghost cells allowed us to follow individual filaments through different heights of the cell and show that the filaments can undergo 90° turns within a thickness of <300 nm (Figure 4). Interestingly, other IFs such as vimentin might not be as flexible, as they seem to be fluctuating less through the different heights of the ghost cells (Figure 4–figure supplement 1).

The bending property of keratin filaments has enabled us to analyze cross section views and therefore to directly reveal that they are built from 6 sub-filaments surrounding an electron dense core. This suggests, that these filaments are composed of 6 tetrameric protofilaments, yielding 24 polypeptides in cross section. In support of this finding, previous mass-per-unit-length studies identified 21 – 25 polypeptides per cross section of reassembled epidermal keratin extracts or in recombinant K8/K18 filaments prepared *in vitro* (28, 29). It is unlikely that the identified sub-filaments represent protofibrils, i.e. octameric assemblies, as this would yield 48 polypeptides in cross section, which does not agree with previous studies. However, MPL studies also found filament populations with a slightly higher or lower numbers of subunits, indicating a polymorphic composition of KIFs. Our data support this, as different 2D classes and raw cross-sectional views reveal a divergence from the hexameric arrangement that would potentially accommodate the previously reported polymorphism.

It is noteworthy that the keratin filaments are not completely filled, but possess a distinct density in the center that is separated from the protofilaments forming the filament tube. This is very similar to the trichocyte keratins found in wool, where an internal core was also identified (30, 64, 65). This seems to be a feature that epidermal and trichocyte keratins have in common, although their amino acid sequences, as well as their arrangement in cells and their functions differ. The internal density may correspond to an additional protofilament (30) or another cellular component.

Polymorphism, i.e. a variable number of protofilaments composing keratin filaments, might also explain the alterations of the helical pattern seen in the 2D class averages (28, 29, 66). Structural flexibility and changes in helical packing of the filament may provide structural support for the elastic nature of the keratin network and would help to explain their high resistance to breakage (4, 46). This feature of keratin filaments would coincide with their task to adapt to different mechanical stresses while maintaining a stable network. Our results also show large heterogeneities in the filament thickness using 2D classification and filament reconstitution methods. Although the most prevalent diameter detected in keratin filaments is 10.1 nm, the diameter can fluctuate between 9.2 – 13.2 nm. This type of heterogeneity has been previously described for several types of IFs (28, 29, 66) and is thought to reflect a varying number of subunits per cross section. However, our results show that regions of increased diameter frequently yield insights into the subunit organization of the filament, indicating that the individual protofilaments are more loosely packed. The increased widths may therefore also reflect regions where rearrangements of the filament take place, subunits might be exchanged (67-70) or interactions with keratin binding proteins might occur. Further, they might reflect areas of local post-translational modifications or regions where filaments were locally damaged. Interestingly, these regions are not restricted to the edges of filaments, but can occur in the mid regions of already assembled filaments. Filament stretches with diameters smaller than 10 nm might reflect either supercoiling or regions that have experienced a greater degree of localized mechanical stress. Previous studies showed that upon stretching beyond 200 %, keratins adopt a plastic behavior that is accompanied by strain hardening and a significant reduction in diameter (46, 71, 72), which is thought to be mainly a result of α-helix to β-sheet transitions of the coiled-coil domains (71-73). Our findings that there are thinner regions in keratin filaments may reflect local domains where extensive forces were applied to the filament network and the filaments adapted by unfolding their coiled-coil domains, leading to a reduction in diameter.

Overall, our results demonstrate that structural polymorphism is an intrinsic property of keratin filaments that were assembled within epithelial cells. Especially when compared to the other components of the cytoskeleton, microtubules and actin filaments, which adapt a highly uniform arrangement, the structural heterogeneity of keratin filaments is clearly exceptional.

Our findings demonstrate the importance of determining a high-resolution structure of keratin filaments in order to understand the details of their assembly states and the functional significance of their heterogeneity in cells. Analyzing *in vivo* assembled filaments provides an approach to study the structure of keratins in their native state with their original post-translational modifications. Understanding the high-resolution structure of keratin filaments would also provide a foundation for determining how keratin mutations affect their structure and how they interact with binding partners. Cryo-EM and cryo-ET are the methods of choice for unravelling the complexities of the 3D structure of mature keratin filaments, as the coordinated use of these techniques can resolve both their flexibility and heterogeneity. Given the rapid advances in cryo-EM imaging, sample preparation and image processing, we anticipate that the structural analysis of keratin intermediate filaments will continue to provide new insights into their cellular structure and functions.

## Material and Methods

### Generation of the K5/K14_1 cell line

Mouse keratinocytes lacking the entire type I keratin cluster (KtyI^-/-^) but stably transfected with K14 (38) were cultured on Collagen I (bovine, CellSystems) coated dishes at 32 °C and 5% CO_2_ in calcium-depleted FAD medium in the presence of puromycin (LabForce, 8 µg/ml medium). Confluent cells were trypsinized using 2.5 x trypsin/EDTA solution (Sigma-Aldrich, T4174) and re-seeded at a maximum splitting ratio of 1:2. To knock-out the K6a and K6b genes, cells were transfected with the pX458 (pSpCas9(BB)-2A-GFP) plasmid (Addgene) carrying a GFP-tagged Cas9 and a guideRNA insert targeting both the K6a and K6b gene (guideRNA sequence: GAGCCACCGCTGCCCCGGGAG). Transfection using electroporation was performed according to the manufacturer’s protocol using a P3 primary cell 4D-Nucleofector kit (Lonza) and program 138 for human keratinocytes, followed by another round of transfection using jetPRIME (Polyplus transfection). Next, genomic DNA was extracted from clonal cell lines using the GenElute Mammalian Genomic DNA kit (Sigma-Aldrich) and the K6a and K6b gene fractions where the indel mutations were expected were amplified by PCR. PCR fragments were sequenced (Microsynth) and the indel mutation spectrum was analyzed using the TIDE webtool (https://tide.nki.nl/). The K5/K14_1 clone was identified as homogenous K6a knockout and heterogenous K6b knockout and was therefore used for all studies. To further verify the knockout, PCR fragments were cloned into the pGEM-T Easy vector (Promega, A1360) and amplified in DH5α cells. Bacterial clones carrying individual gene sequences of K6a or K6b were picked and amplified, plasmids were extracted and sequenced (Microsynth). By analyzing 19 K6a and 22 K6b sequences, the homogenous K6a and heterogenous K6b knockout were verified.

### Immunostaining

The KtyI^-/-^ K14 and K5/K14_1 cells were seeded on Collagen I coated glass cover slips in cell culture dishes and incubated overnight at 32 °C and in 5% CO_2_. For staining with keratin antibodies, cells were fixed for 5 min using ice-cold 99.9% anhydrous methanol (Alfa Aesar, 41838). Non-specific antibody binding sites were blocked by incubating the cover slips for 30 min in blocking buffer (1% BSA, 22.52 mg/ml glycine in PBS with 0.1% Tween (PBS-T)). Next, cover slips were incubated for 1 h at room temperature with mouse anti-mouse Keratin 14 (LL02, Thermo Fisher, MA5-11599, 1:100 – 1:10), rabbit anti-mouse Keratin 5 (BioLegend, 905503, 1:500) or rabbit anti-mouse Keratin 6a (BioLegend, 905702, 1:500) in 1% BSA in 0.1% PBS-T. It should be noted here that the K6a antibody used is a polyclonal antibody, which is suspected to bind to the K6b protein as well, due to their high sequence identity (92.6%). After 3x 5 min washing steps in PBS, cover slips were incubated with Cy3 donkey anti-rabbit (Jackson Immuno Research, 711-165-152, 1:100) or FITC donkey anti-mouse (Jackson Immuno Research, 715-095-150, 1:100) secondary antibodies in 1% BSA in 0.1% PBS-T. Cells were washed 3x for 5 min in PBS, before nuclei were stained with Hoechst 33342 (Sigma-Aldrich, B2261, 1:10,000) for 10 – 20 min at room temperature. After a final wash step 3x for 5 min in PBS, the cover slips were mounted on glass slides with Dako mounting medium (Agilent, S3023) or Prolong Glass Anti-Fade (Thermo Fisher, P36980). Confocal imaging for Figure 1 was carried out with a laser scanning confocal microscope (Nikon A1R confocal microscope, Nikon) using an oil immersion objective lens (Plan Apo 60X Oil objective, 1.4 NA, Nikon). Keratin was excited with a 488 nm wavelength laser and the optical sections were imaged at 100 nm intervals. Maximum intensity projections of the Z-stacks are presented. Keratin networks for Figure 1–figure supplement 1 were imaged using a spinning disk confocal laser scanning microscope (Olympus IXplore SpinSR10 with YOKOGAWA CSU-W1 spinning disk). 3D confocal stacks were acquired with a UPLSAPO UPlan S Apo 60x/1.3 OIL objective (Olympus). Fluorescent proteins were excited at 405 nm (50 mW, 10 % laser power), 488 nm (100 mW, 15 % laser power) and 561 nm (100 mW, 5 % laser power).

### Sample preparation for cryo-EM and cryo-ET

K5/K14_1 cells were seeded on glow-discharged Collagen I coated holey carbon gold EM grids (Au R2/1, 200 mesh, Quantifoil) and incubated overnight at 32 °C and in 5% CO_2_. The grids were rinsed in washing buffer (1x PBS, 2 mM MgCl_2_), cells were permeabilized for 15 – 20 s in permeabilization buffer (1x PBS, 0.1% Triton X-100, 600 mM KCl, 10 mM MgCl_2_ and protease inhibitors), and rinsed again in PBS. Next, the grids were incubated with 2.5 units/µl benzonase (Merck, 71206-3) in washing buffer for 30 min and washed again before vitrification in liquid ethane using a manual plunge freezing device. For cryo-ET samples, 10 nm gold fiducial markers (Aurion, Netherlands) were added to the grids right before freezing.

### Cryo-EM and cryo-ET data acquisition

The grids were analyzed using a 300 kV Titan Krios electron microscope (Thermo Fisher) equipped with a K2 Summit direct electron detector (Gatan) mounted on a post-column energy filter (Gatan). Cryo-EM micrographs were acquired in zero-loss energy mode using a 20 eV slit. Data were recorded with SerialEM 3.5.8 in low dose mode (74). Micrographs were acquired at nominal magnifications of 46,511 x with a pixel size of 1.075 Å, 28,571 x with a pixel size of 1.75 Å and 22,665 x with a pixel size of 2.206 Å. A defocus range between -0.5 and -3.5 μm was chosen. Dose-fractionation was used with a frame exposure of 0.2 s with a total exposure time of 10 s (50 frames in total). This corresponds to a total electron dose of ∼20 e/Å^2^ for the 22,665 x dataset, ∼41 e/Å^2^ for the 28,571 x and ∼84 e/Å^2^ for the 46,511 x dataset.

Tilt series were acquired in zero-loss energy mode with a 20 eV slit at a nominal magnification of 28,571 x, resulting in a pixel size of 1.75 Å and a defocus of -3 µm. A bidirectional tilt scheme with a tilt range of ± 60° and an increment of 3° was chosen, corresponding to 41 projections per tilt series and a total accumulative electron dose of ∼89 e/Å^2^. SerialEM 3.5.8 in low dose mode was used for data acquisition (74).

### Minimal apparent persistence length measurements

Highly bent keratin filaments were identified in electron micrographs and traced with Fiji using the segmented line tool including spline fit. Minimal filament contour lengths that undergo a 90° turn were traced. The persistence length is defined as the distance along a filament at which the tangent-tangent correlation function along the contour length decays, this occurs after a 90° turn (75). However, since our sample is out of equilibrium, as individual filaments are entangled in a network and absorbed to the EM grid, and filaments are imaged in 2D, only an apparent persistence length is described. Further, only highly bent filaments were considered in this analysis, yielding a minimal apparent persistence length, as the whole filament population is diverse and cannot be described as a single state.

### Cryo-EM data processing

1,860 cryo-EM micrographs at a magnification of 22,665x were processed with RELION 2.1 and RELION 3.0 using the helical toolbox (50, 51, 76). Frame-based motion correction and dose-weighting were performed using MotionCor2 (77). The contrast transfer function was estimated using CTFFIND4 (78). Low-quality micrographs showing high defocus, high astigmatism or low resolution were excluded, resulting in 1,763 micrographs used for further processing steps. Keratin filaments were either picked manually or automatically using the RELION helical toolbox. To generate a template for autopicking, 55,073 keratin particles were picked manually as start-to-end helices, extracted with a box size of 250 pixels (∼55 nm) and 2D classified twice to create classes with straight keratin segments. These classes served as a reference for automated picking of 505,211 particles. For manual picking, 298,056 particles were selected as start-to-end helices. Particles were extracted in boxes of 250 pixels, corresponding to ∼55 nm, or 164 pix, corresponding to ∼36 nm, with an inter-box distance of 50 Å. Iterative 2D classification procedures were performed, using a spherical mask of 500 Å or 356 Å, respectively.

Keratin filament segments, 55 nm in length, were classified to yield 305,495 particles in straight classes. Autocorrelation spectra were calculated with MATLAB (2019a, MathWorks). The filament diameter was measured by plotting intensity line-profiles of all classes using MATLAB and measuring the area where the intensity lies above zero. OriginPro 2018 software (OriginLab Corporation) was used to plot the diameter distribution. Intensity line-profiles of each class were generated in MATLAB by averaging all lateral sections through the segment. A mean intensity line-profile was generated by averaging all classes of similar diameter (Figure 2–figure supplement 1A).

Segments with a box size of 36 nm were used for *in silico* filament reconstitution. Filament reconstitution was performed as previously reported (41) and described below with classes of automatically, as well as manually, picked particles.

Actin filaments were processed identically to keratin filaments to guarantee comparability. To generate a template for autopicking, 22,228 actin particles were picked manually as start-to-end helices, extracted with a box size of 164 pixels (∼36 nm) and 2D classified twice to create classes with straight actin segments. These classes served as a reference for automated picking of 693,903 particles. Particles were extracted in boxes of 164 pixels, corresponding to ∼36 nm, with an inter-box distance of 50 Å. Multiple rounds of 2D classification were performed, using a spherical mask of 356 Å. 405,044 particles from the highest resolved 2D classes were used for 3D classification into 5 classes. The highest resolved 3D class, containing 174,954 particles, was projected to 3D refinement. The final unmasked map showed a resolution of 7.38 Å, based on the gold standard Fourier shell correlation (FSC) 0.143 criterion (50, 79). The structure was sharpened to 6.13 Å using an isotropic B-factor of -276 Å^2^.

### Cryo-ET data processing

Tilt series were processed using the IMOD workflow, including contrast transfer function (CTF) correction (80). For visualization purposes, a SIRT-like filter according to 10 iterations was applied during tomogram reconstruction. Cellular structures present in the tomograms were manually segmented and visualized using the Amira 5.6.0 software package (Thermo Fischer Scientific). 710 cross section views of keratin filaments were picked in 21 tomograms and reconstructed as sub-tomograms using IMOD. Central 2D slices were extracted from the sub-tomograms and utilized for 2D classification in RELION.

### Reconstitution of keratin filaments

To generate reconstituted filaments a back-mapping strategy was pursued in MATLAB, using the keratin segments which were used for 2D classification. First, all particles belonging to the same filament were grouped. Filament assignments were made based on the helical tube ID defined by RELION for every particle. Next, all particles belonging to the same filament were sorted in ascending order based on their picking coordinates. Then, their corresponding 2D class images were inversely transformed, so that their orientation matches the original orientation of the raw segments in the cryo-EM micrographs. Next, the 2D class images were plotted at the original coordinates of the particles. To remove background noise, the classes were masked in the y-direction and only the central 132 Å were plotted. Since particles were picked with inter-box distances of 50 Å, while 2D classes have a box size of 360 Å, neighboring segments would strongly overlap. To avoid this, classes were cropped to not extend into neighboring particle positions, and only a small amount of overlap of <4 with soft edges was allowed to avoid cropped edges in slightly bent filaments. Reconstituted filaments were normalized to equal intensity. Next, a straightening procedure was applied as previously described to extract, align and straighten the reconstituted filaments (81, 82). Validity of this approach was ensured by using high-resolution actin classes as a control.

### Analyzing helical and straight-line patterns in individual filaments

Plots representing the order of helical and straight segments along individual filaments, represented by colored circles, were generated in MATLAB as previously described (53). 2D classes were grouped into helical or straight clusters based on their appearance. Next, each particle within a filament was represented by red or blue circles, depending on whether its corresponding 2D class belonged to the helical or straight cluster. In the analysis seen in Figure 3–figure supplement 1B, segments that originate from the same filament are plotted as columns of circles. Segments are sorted in ascending order based on their coordinates along the filament.

### 3D reconstruction of a keratin filament

305,495 uniform keratin segments from 55 nm boxes were selected by 2D classification and used for 3D reconstruction. To generate a low-resolution 3D filament model, the rotation angle along the filament axis of all particles was randomized to prevent preferred orientations. Next, a filament was reconstructed using relion_reconstruct. The 3D model was visualized using Chimera (83).

## Data availability

Representative cryo-ET data have been deposited in the Electron Microscopy Data Bank under accession code xxx.

## Acknowledgements

This research was funded by the Swiss National Science Foundation Grant (31003A, 179418). M.S.W. was supported by the Forschungskredit of the University of Zurich [FK-18-041]. The Goldman laboratory is supported by grants 5PO1 GM096971 and RO1GM140108 from the National Institutes of Health. The authors would like to thank the Center for Microscopy and Image Analysis at the University of Zurich for providing support and equipment.

## Author contributions

M. S. W. performed the CRISPR/Cas9 knockout, prepared samples, recorded and analyzed data.

M. S. W and M. E. developed methods. M. S. W. and S. S. acquired fluorescent images. T. M. M. provided the KtyI^-/-^ K14 cell line and reviewed the manuscript. O. M. provided resources, funding and administration. M. S. W. together with R. D. G. and O. M. conceived the research and wrote the manuscript.

## Declaration of interests

The authors declare no conflicts of interest.

## Notes

### Competing Interest Statement

The authors have declared no competing interest.

## References

1. Szeverenyi I, Cassidy AJ, Chung CW, Lee BT, Common JE, Ogg SC, et al. The Human Intermediate Filament Database: comprehensive information on a gene family involved in many human diseases. Hum Mutat. 2008;29(3):351–60.

2. Bragulla HH, Homberger DG. Structure and functions of keratin proteins in simple, stratified, keratinized and cornified epithelia. J Anat. 2009;214(4):516–59.

3. Toivola DM, Boor P, Alam C, Strnad P. Keratins in health and disease. Current opinion in cell biology. 2015;32:73–81.

4. Etienne-Manneville S. Cytoplasmic Intermediate Filaments in Cell Biology. Annu Rev Cell Dev Biol. 2018;34:1–28.

5. Pora A, Yoon S, Dreissen G, Hoffmann B, Merkel R, Windoffer R, et al. Regulation of keratin network dynamics by the mechanical properties of the environment in migrating cells. Scientific reports. 2020;10(1):4574.

6. Kolsch A, Windoffer R, Wurflinger T, Aach T, Leube RE. The keratin-filament cycle of assembly and disassembly. Journal of cell science. 2010;123(Pt 13):2266–72.

7. Robert A, Hookway C, Gelfand VI. Intermediate filament dynamics: What we can see now and why it matters. sBioessays. 2016;38(3):232–43.

8. Windoffer R, Woll S, Strnad P, Leube RE. Identification of novel principles of keratin filament network turnover in living cells. Molecular biology of the cell. 2004;15(5):2436–48.

9. Yoon KH, Yoon M, Moir RD, Khuon S, Flitney FW, Goldman RD. Insights into the dynamic properties of keratin intermediate filaments in living epithelial cells. Journal of Cell Biology. 2001;153(3):503–16.

10. Haines RL, Lane EB. Keratins and disease at a glance. J Cell Sci. 2012;125(Pt 17):3923–8.

11. Jacob JT, Coulombe PA, Kwan R, Omary MB. Types I and II Keratin Intermediate Filaments. Cold Spring Harbor perspectives in biology. 2018;10(4).

12. Coulombe PA, Hutton ME, Letai A, Hebert A, Paller AS, Fuchs E. Point mutations in human keratin 14 genes of epidermolysis bullosa simplex patients: genetic and functional analyses. Cell. 1991;66(6):1301–11.

13. Coulombe PA, Lee CH. Defining keratin protein function in skin epithelia: epidermolysis bullosa simplex and its aftermath. J Invest Dermatol. 2012;132(3 Pt 2):763–75.

14. Wilson AK, Coulombe PA, Fuchs E. The roles of K5 and K14 head, tail, and R/K L L E G E domains in keratin filament assembly in vitro. The Journal of cell biology. 1992;119(2):401–14.

15. Parry DA, Steinert PM. Intermediate filaments: molecular architecture, assembly, dynamics and polymorphism. Q Rev Biophys. 1999;32(2):99–187.

16. Chernyatina AA, Guzenko D, Strelkov SV. Intermediate filament structure: the bottom-up approach. Current opinion in cell biology. 2015;32:65–72.

17. Steinert PM, Marekov LN, Fraser RD, Parry DA. Keratin intermediate filament structure. Crosslinking studies yield quantitative information on molecular dimensions and mechanism of assembly. Journal of molecular biology. 1993;230(2):436–52.

18. Lee CH, Kim MS, Li S, Leahy DJ, Coulombe PA. Structure-Function Analyses of a Keratin Heterotypic Complex Identify Specific Keratin Regions Involved in Intermediate Filament Assembly. Structure. 2020;28(3):355–62 e4.

19. Herrmann H, Aebi U. Intermediate Filaments: Structure and Assembly. Cold Spring Harbor perspectives in biology. 2016;8(11).

20. Eldirany SA, Lomakin IB, Ho M, Bunick CG. Recent insight into intermediate filament structure. Current opinion in cell biology. 2020;68:132–43.

21. Lee CH, Kim MS, Chung BM, Leahy DJ, Coulombe PA. Structural basis for heteromeric assembly and perinuclear organization of keratin filaments. Nat Struct Mol Biol. 2012;19(7):707-+.

22. Bunick CG, Milstone LM. The X-Ray Crystal Structure of the Keratin 1-Keratin 10 Helix 2B Heterodimer Reveals Molecular Surface Properties and Biochemical Insights into Human Skin Disease. J Invest Dermatol. 2017;137(1):142–50.

23. Eldirany SA, Ho M, Hinbest AJ, Lomakin IB, Bunick CG. Human keratin 1/10-1B tetramer structures reveal a knob-pocket mechanism in intermediate filament assembly. The EMBO journal. 2019;38(11).

24. Bray DJ, Walsh TR, Noro MG, Notman R. Complete Structure of an Epithelial Keratin Dimer: Implications for Intermediate Filament Assembly. PloS one. 2015;10(7):e0132706.

25. Astbury WT. X-Ray Studies of the Structure of Compounds of Biological Interest. Annual review of biochemistry. 1939;8(1):113–33.

26. Crick FHC. Is Alpha-Keratin a Coiled Coil. Nature. 1952;170(4334):882–3.

27. Crick FHC. The Packing of Alpha-Helices - Simple Coiled-Coils. Acta Crystallogr. 1953;6(8-9):689–97.

28. Herrmann H, Haner M, Brettel M, Ku NO, Aebi U. Characterization of distinct early assembly units of different intermediate filament proteins. Journal of molecular biology. 1999;286(5):1403–20.

29. Engel A, Eichner R, Aebi U. Polymorphism of Reconstituted Human Epidermal Keratin Filaments - Determination of Their Mass-Per-Length and Width by Scanning-Transmission Electron-Microscopy (Stem). J Ultra Mol Struct R. 1985;90(3):323–35.

30. Bruce Fraser RD, Steinert PM, Parry DAD. Structural changes in trichocyte keratin intermediate filaments during keratinization. Journal of structural biology. 2003;142(2):266–71.

31. Parry DA. Hard alpha-keratin intermediate filaments: an alternative interpretation of the low-angle equatorial X-ray diffraction pattern, and the axial disposition of putative disulphide bonds in the intra-and inter-protofilamentous networks. Int J Biol Macromol. 1996;19(1):45–50.

32. Fraser RDB, Parry DAD. Intermediate filament structure in fully differentiated (oxidised) trichocyte keratin. Journal of structural biology. 2017;200(1):45–53.

33. Watts NR, Jones LN, Cheng NQ, Wall JS, Parry DAD, Steven AC. Cryo-electron microscopy of trichocyte (hard alpha-keratin) intermediate filaments reveals a low-density core. Journal of structural biology. 2002;137(1-2):109–18.

34. Herrmann H, Aebi U. Intermediate filaments: Molecular structure, assembly mechanism, and integration into functionally distinct intracellular scaffolds. Annual review of biochemistry. 2004;73:749–89.

35. Weber MS, Wojtynek M, Medalia O. Cellular and Structural Studies of Eukaryotic Cells by Cryo-Electron Tomography. Cells. 2019;8(1).

36. Scheres SH. Processing of Structurally Heterogeneous Cryo-EM Data in RELION. Methods Enzymol. 2016;579:125–57.

37. Moll R, Divo M, Langbein L. The human keratins: biology and pathology. Histochemistry and Cell Biology. 2008;129(6):705–33.

38. Homberg M, Ramms L, Schwarz N, Dreissen G, Leube RE, Merkel R, et al. Distinct Impact of Two Keratin Mutations Causing Epidermolysis Bullosa Simplex on Keratinocyte Adhesion and Stiffness. J Invest Dermatol. 2015;135(10):2437–45.

39. Hu J, Li Y, Hao Y, Zheng T, Gupta SK, Parada GA, et al. High stretchability, strength, and toughness of living cells enabled by hyperelastic vimentin intermediate filaments. Proceedings of the National Academy of Sciences of the United States of America. 2019;116(35):17175–80.

40. Turgay Y, Eibauer M, Goldman AE, Shimi T, Khayat M, Ben-Harush K, et al. The molecular architecture of lamins in somatic cells. Nature. 2017;543(7644):261–4.

41. Kronenberg-Tenga R, Tatli M, Eibauer M, Wu W, Shin JY, Bonne G, et al. A lamin A/C variant causing striated muscle disease provides insights into filament organization. Journal of cell science. 2021.

42. Svitkina TM, Borisy GG. Correlative light and electron microscopy of the cytoskeleton of cultured cells. Methods Enzymol. 1998;298:570–92.

43. Svitkina TM, Verkhovsky AB, Borisy GG. Improved procedures for electron microscopic visualization of the cytoskeleton of cultured cells. Journal of structural biology. 1995;115(3):290–303.

44. Sailer M, Hohn K, Luck S, Schmidt V, Beil M, Walther P. Novel electron tomographic methods to study the morphology of keratin filament networks. Microsc Microanal. 2010;16(4):462–71.

45. Tenga R, Medalia O. Structure and unique mechanical aspects of nuclear lamin filaments. Curr Opin Struct Biol. 2020;64:152–9.

46. Block J, Schroeder V, Pawelzyk P, Willenbacher N, Koster S. Physical properties of cytoplasmic intermediate filaments. Biochim Biophys Acta. 2015;1853(11 Pt B):3053–64.

47. Coulombe PA, Fuchs E. Elucidating the early stages of keratin filament assembly. The Journal of cell biology. 1990;111(1):153–69.

48. Herrmann H, Wedig T, Porter RM, Lane EB, Aebi U. Characterization of early assembly intermediates of recombinant human keratins. Journal of structural biology. 2002;137(1-2):82–96.

49. Koster S, Weitz DA, Goldman RD, Aebi U, Herrmann H. Intermediate filament mechanics in vitro and in the cell: from coiled coils to filaments, fibers and networks. Current opinion in cell biology. 2015;32:82–91.

50. Scheres SHW. RELION: Implementation of a Bayesian approach to cryo-EM structure determination. Journal of structural biology. 2012;180(3):519–30.

51. He SD, Scheres SHW. Helical reconstruction in RELION. Journal of structural biology. 2017;198(3):163–76.

52. Diaz R, Rice WJ, Stokes DL. Fourier-Bessel reconstruction of helical assemblies. Methods Enzymol. 2010;482:131–65.

53. Martins B, Sorrentino S, Chung WL, Tatli M, Medalia O, Eibauer M. Unveiling the polarity of actin filaments by cryo-electron tomography. Structure. 2021.

54. Redmond CJ, Coulombe PA. Intermediate filaments as effectors of differentiation. Curr Opin Cell Biol. 2021;68:155–62.

55. Gordon BA. Neurofilaments in disease: what do we know? Curr Opin Neurobiol. 2020;61:105–15.

56. Danielsson F, Peterson MK, Caldeira Araujo H, Lautenschlager F, Gad AKB. Vimentin Diversity in Health and Disease. Cells. 2018;7(10).

57. Snider NT, Omary MB. Post-translational modifications of intermediate filament proteins: mechanisms and functions. Nature reviews Molecular cell biology. 2014;15(3):163–77.

58. Sigworth FJ. Principles of cryo-EM single-particle image processing. Microscopy (Oxf). 2016;65(1):57–67.

59. Quinlan RA, Cohlberg JA, Schiller DL, Hatzfeld M, Franke WW. Heterotypic tetramer (A2D2) complexes of non-epidermal keratins isolated from cytoskeletons of rat hepatocytes and hepatoma cells. Journal of molecular biology. 1984;178(2):365–88.

60. Gittes F, Mickey B, Nettleton J, Howard J. Flexural rigidity of microtubules and actin filaments measured from thermal fluctuations in shape. The Journal of cell biology. 1993;120(4):923–34.

61. Sapra KT, Qin Z, Dubrovsky-Gaupp A, Aebi U, Muller DJ, Buehler MJ, et al. Nonlinear mechanics of lamin filaments and the meshwork topology build an emergent nuclear lamina. Nature communications. 2020;11(1):6205.

62. Lichtenstern T, Mucke N, Aebi U, Mauermann M, Herrmann H. Complex formation and kinetics of filament assembly exhibited by the simple epithelial keratins K8 and K18. Journal of structural biology. 2012;177(1):54–62.

63. Pawelzyk P, Mucke N, Herrmann H, Willenbacher N. Attractive Interactions among Intermediate Filaments Determine Network Mechanics In Vitro. PloS one. 2014;9(4).

64. Rogers GE. Structural and biochemical features of the hair follicle. Academic Press. 1964(The Epidermis):179–236.

65. Kadir M, Wang XW, Zhu BW, Liu J, Harland D, Popescu C. The structure of the “amorphous” matrix of keratins. Journal of structural biology. 2017;198(2):116–23.

66. Steven AC, Hainfeld JF, Trus BL, Wall JS, Steinert PM. The distribution of mass in heteropolymer intermediate filaments assembled in vitro. Stem analysis of vimentin/desmin and bovine epidermal keratin. The Journal of biological chemistry. 1983;258(13):8323–9.

67. Colakoglu G, Brown A. Intermediate filaments exchange subunits along their length and elongate by end-to-end annealing. Journal of Cell Biology. 2009;185(5):769–77.

68. Ngai J, Coleman TR, Lazarides E. Localization of newly synthesized vimentin subunits reveals a novel mechanism of intermediate filament assembly. Cell. 1990;60(3):415–27.

69. Miller RK, Vikstrom K, Goldman RD. Keratin Incorporation into Intermediate Filament Networks Is a Rapid Process. Journal of Cell Biology. 1991;113(4):843–55.

70. Vikstrom KL, Borisy GG, Goldman RD. Dynamic aspects of intermediate filament networks in BHK-21 cells. Proceedings of the National Academy of Sciences of the United States of America. 1989;86(2):549–53.

71. Kreplak L, Bar H, Leterrier JF, Herrmann H, Aebi U. Exploring the mechanical behavior of single intermediate filaments. Journal of molecular biology. 2005;354(3):569–77.

72. Fudge DS, Gardner KH, Forsyth VT, Riekel C, Gosline JM. The mechanical properties of hydrated intermediate filaments: insights from hagfish slime threads. Biophys J. 2003;85(3):2015–27.

73. Pinto N, Yang FC, Negishi A, Rheinstadter MC, Gillis TE, Fudge DS. Self-assembly enhances the strength of fibers made from vimentin intermediate filament proteins. Biomacromolecules. 2014;15(2):574–81.

74. Mastronarde DN. Automated electron microscope tomography using robust prediction of specimen movements. Journal of structural biology. 2005;152(1):36–51.

75. Reisner W, Pedersen JN, Austin RH. DNA confinement in nanochannels: physics and biological applications. Rep Prog Phys. 2012;75(10):106601.

76. Zivanov J, Nakane T, Forsberg BO, Kimanius D, Hagen WJH, Lindahl E, et al. New tools for automated high-resolution cryo-EM structure determination in RELION-3. Elife. 2018;7.

77. Zheng SQ, Palovcak E, Armache JP, Verba KA, Cheng YF, Agard DA. MotionCor2: anisotropic correction of beam-induced motion for improved cryo-electron microscopy. Nature methods. 2017;14(4):331–2.

78. Rohou A, Grigorieff N. CTFFIND4: Fast and accurate defocus estimation from electron micrographs. Journal of structural biology. 2015;192(2):216–21.

79. Rosenthal PB, Henderson R. Optimal determination of particle orientation, absolute hand, and contrast loss in single-particle electron cryomicroscopy. Journal of molecular biology. 2003;333(4):721–45.

80. Kremer JR, Mastronarde DN, McIntosh JR. Computer visualization of three-dimensional image data using IMOD. Journal of Structural Biology. 1996;116(1):71–6.

81. Steinert PM, Steven AC, Roop DR. The molecular biology of intermediate filaments. Cell. 1985;42(2):411–20.

82. Kocsis E, Trus BL, Steer CJ, Bisher ME, Steven AC. Image averaging of flexible fibrous macromolecules: the clathrin triskelion has an elastic proximal segment. Journal of structural biology. 1991;107(1):6–14.

83. Pettersen EF, Goddard TD, Huang CC, Couch GS, Greenblatt DM, Meng EC, et al. UCSF Chimera--a visualization system for exploratory research and analysis. J Comput Chem. 2004;25(13):1605–12.

