## Supplementary figures for "Structural heterogeneity of cellular K5/K14 filaments as revealed by cryo-electron microscopy"

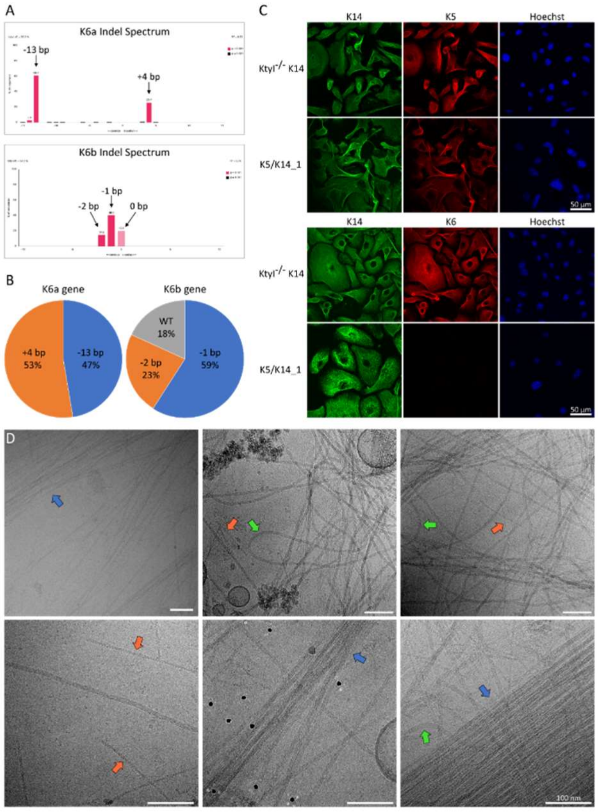


**Figure 1–figure supplement 1. Knockout of K6 isoforms by CRISPR/Cas9.**

(A) TIDE analysis of the K6a and K6b gene sequences of the K5/K14_1 cell line carrying the mutations induced by non-homologous end joining. Genomic DNA fragments of the K6a and K6b gene were amplified by PCR and sequenced. Peaks in the Indel spectrum confirm mis-sense insertion or deletion mutations in each gene. For K6a, a deletion of 13 base pairs (bp) and an insertion of 4 bp were detected, while for the K6b gene a deletion of 2 bp and 1 bp were detected, as well as the wildtype sequence. (B) Pie chart plot of the frequency of mutations that were detected in the K6a and K6b gene of the K5/K14_1 cell line. For this analysis, PCR amplified fragments of genomic K6a and K6b DNA were ligated into the pGEM T-Easy vector and transformed into bacteria. Bacterial clones which took up individual plasmids carrying a specific mutation were cultivated and the plasmids were extracted individually and analyzed. For the K6a gene, 19 bacterial clones were analyzed, for the K6b gene, 22 bacterial clones were analyzed. (C) Immunofluorescence analysis of Ktyl$^{-/-}$ K14 cells and K5/K14_1 cells co-stained for K5 (red) and K14 (green) or K6 (red) and K14 (green). DNA was labelled with Hoechst 33342 (blue). No filamentous K6 network could be detected in the K5/K14_1 cells. Scale bars: 50 µm. (D) Representative cryo-EM micrographs of the K5/K14 network in K5/K14_1 ghost cells. Keratin filaments can be clearly distinguished from actin filaments (orange arrows). Keratin bundles (blue arrows) and very wavy filaments (green arrows) are highlighted. Scale bars: 100 nm.

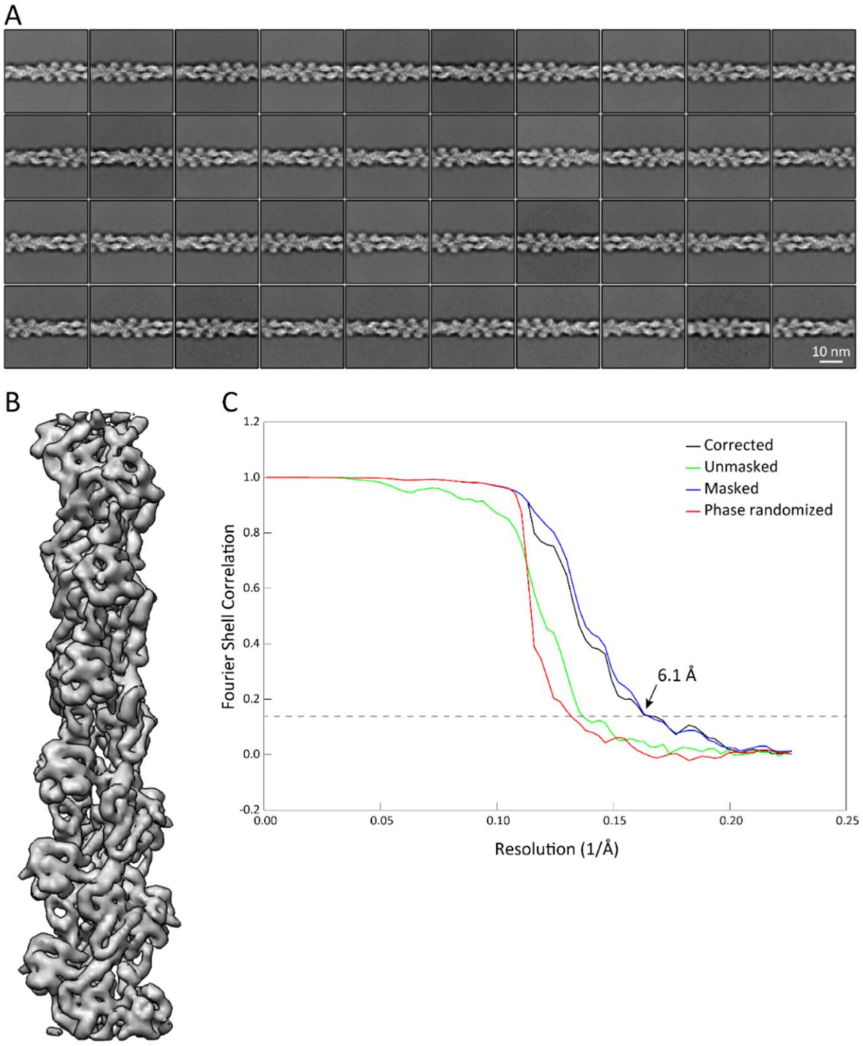


**Figure 1–figure supplement 2. Validation that the sample preparation allows us to retrieve data at sub-nanometer resolution.**

Analysis of actin filaments detected in the K5/K14_1 ghost cells served as internal quality control for the sample preparation and the acquired cryo-EM data. Actin filaments were detected around keratins and analyzed independently. (A) Representative 2D classes of F-actin segments of 36 nm length. (B) 3D refined structure of an *in vivo* assembled actin filament with a resolution of 6.1 Å. Individual α-helices can be recognized. (C) Fourier shell correlation (FSC) plot of the structure shown in (B). The plot shows the unmasked (green), masked (blue), phase randomized (red) and masking-effect corrected (black) FSC curves. The resolution at which the gold-standard FSC curve drops below the 0.143 threshold is indicated.

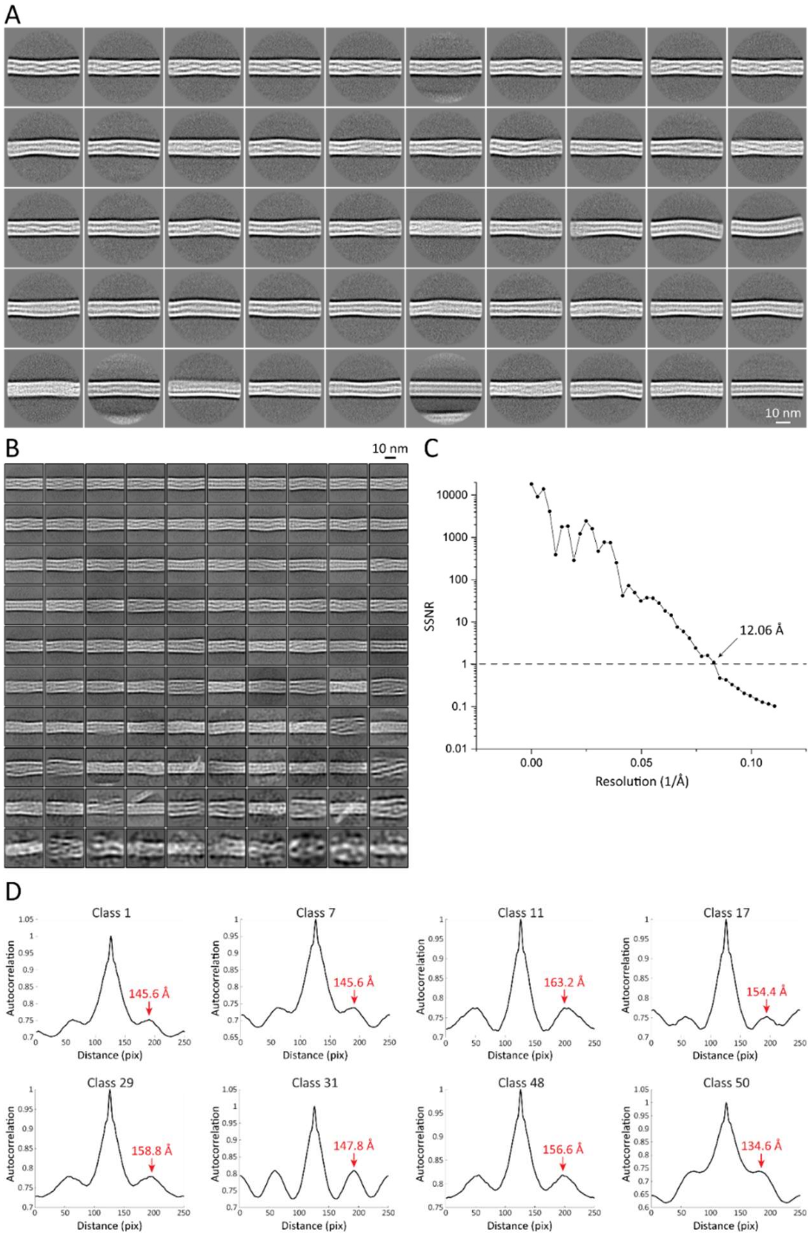


**Figure 2–figure supplement 1. 2D structural analysis of keratin segments.**

(A) Representative 2D class averages of 55 nm long keratin segments. These classes were used for measurements of filament diameter, mean intensity-line profiles and autocorrelation spectra. Particles from these classes were further used to generate the 3D keratin model. (B) Representative 2D class averages of 36 nm long keratin segments. These classes were utilized for reconstitution of long keratin filaments. (C) Plot of the spectral signal-to-noise ratio (SSNR) of the highest resolved 2D class utilized for filament reconstitution. The SSNR was plotted against the resolution. The position before the SSNR drops below 1 indicates the resolution of the 2D class, as until this point the level of signal exceeds the level of noise. The resolution of the class is indicated. (D) Representative autocorrelation spectra of multiple 55 nm long helical keratin classes. Peaks of the autocorrelation function corresponding to the repeating unit in the 2D classes are indicated by labelled red arrows.

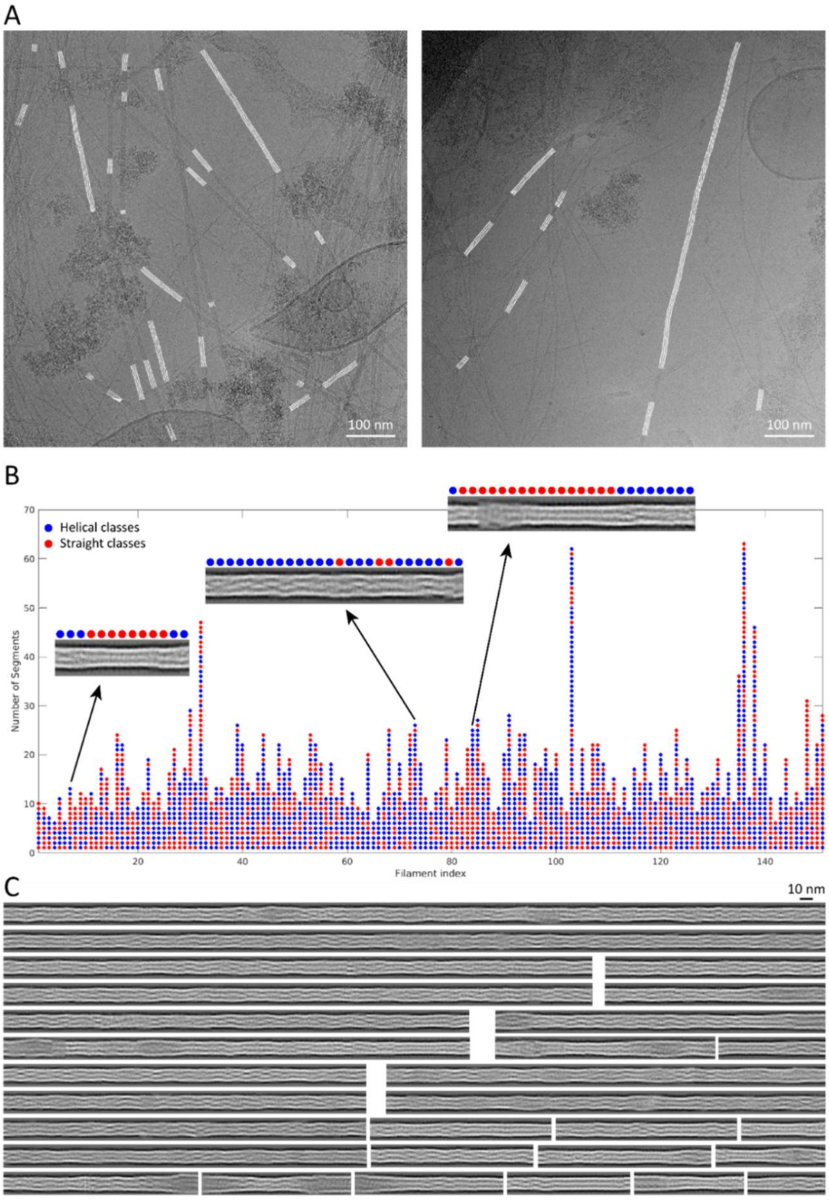


**Figure 3–figure supplement 1. Reconstitution of keratin filaments.**

(A) Back-mapping of class averages onto the original filaments allowed us to assemble reconstituted, non-straightened keratin filaments in the original cryo-EM micrographs. Reconstituted filaments are displayed in white. (B) Keratin segments that originate from the same filament are plotted as columns of circles. The filaments are composed of helical (blue) and straight (red) class averages. Representative reconstituted filaments are shown and their composition by helical and straight classes is indicated. (C) Examples of reconstituted keratin filaments revealing the extensive heterogeneity in filament architecture.

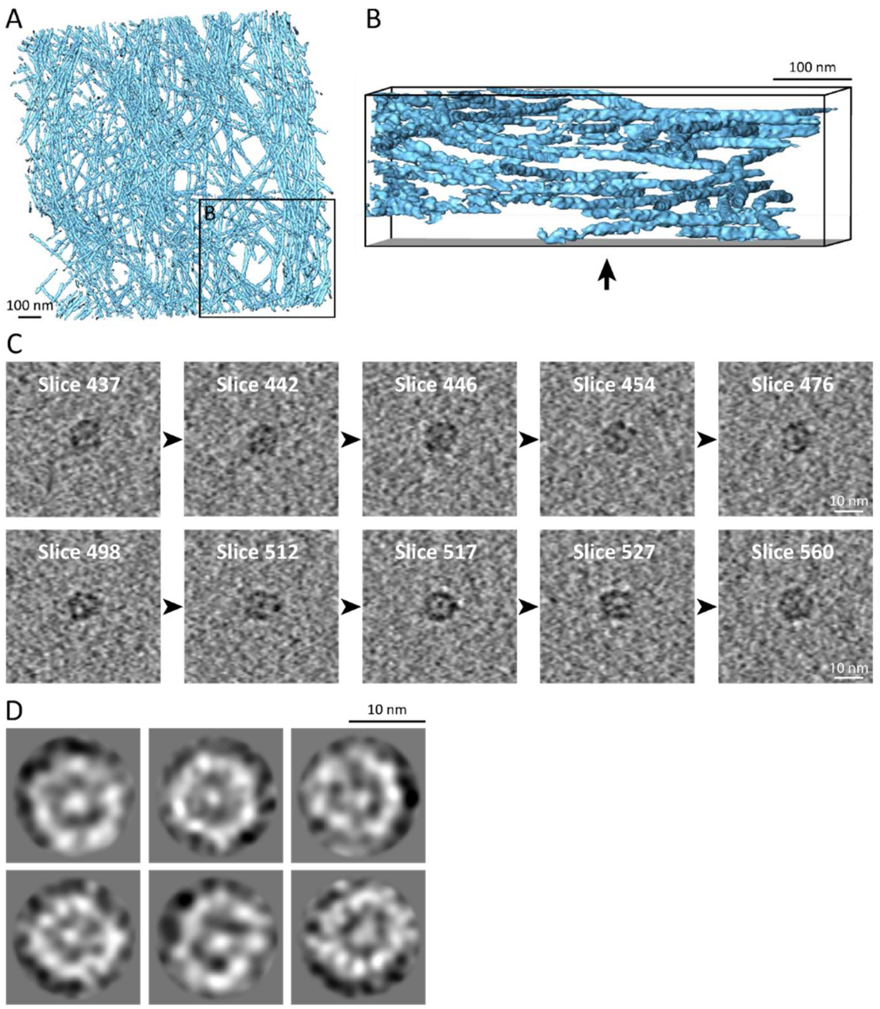


**Figure 4–figure supplement 1. Keratin filaments show a unique flexibility and contain a central density.**

(A) Surface rendering of a tomogram of the vimentin network (blue) in a mouse embryonic fibroblast (MEF) ghost cell. The vimentin network shows reduced flexibility in comparison to keratin filaments, as vimentin filaments span less through the height of the tomographic volume. No cross section views could be detected in 225 vimentin tomograms that were acquired. (B) Rotated view of the area boxed in (A), showing a side view of the vimentin network that reveals less fluctuations through the height of the tomogram volume when compared to keratin filaments (Figure 4C), although the tomograms have a comparable thickness. The arrow indicates the viewing direction from (A). (C) Sequential 7 nm thick xy-slices through two sub-tomograms containing single keratin filaments in cross section views. The top and the bottom row show different filaments. Cutting through these filaments emphasizes the variable appearances detected in different slices and allow the identification of more or less protofilaments, respectively. All slices reveal the internal electron dense core. Scale bars: 10 nm. (D) 2D class averages of CTF-corrected keratin filament cross section views. Individual protofilaments in the ring and an electron dense core can be identified.
